## Supplementary information for "A DCL3 dicing code within Pol IV-RDR2 transcripts diversifies the siRNA pool guiding RNA-directed DNA methylation"

**Supplementary Table 1: RNA oligonucleotides used for DCL3 dicing assays**

| Name | Sequence | Relevant Figure(s) |
|---|---|---|
| **oligos for testing DCL3 dicing of blunt or overhanging ends** | | |
| 37 nt top | 5'- ACAAGCGAAUGAGUCAUUCAUCCUAAGUCCAACAUAG -3' | 1C |
| 37 nt bottom | 5'- CUAUGUUGGACUUAGGAUGAAUGACUCAUUCGCUUGU -3' | 1C |
| 38 nt bottom | 5'- CUAUGUUGGACUUAGGAUGAAUGACUCAUUCGCUUGUA -3' | 1C |
| 39 nt bottom | 5'- CUAUGUUGGACUUAGGAUGAAUGACUCAUUCGCUUGUAG -3' | 1C |
| **oligos for testing DCL3 preferences for overhang length** | | |
| 38 nt top | 5'- ACAAGCGAAUGAGUCAUUCAUCCUAAGUCCAACAUAGU -3' | 1D, 1E, S1B |
| 38 nt bottom | 5'- ACUAUGUUGGACUUAGGAUGAAUGACUCAUUCGCUUGU -3' | 1E |
| 39 nt bottom | 5'- ACUAUGUUGGACUUAGGAUGAAUGACUCAUUCGCUUGUU -3' | 1D, S1B |
| 40 nt bottom | 5'- ACUAUGUUGGACUUAGGAUGAAUGACUCAUUCGCUUGUUU -3' | 1D, S1B |
| **oligos for testing DCL3 cleavage of 24/25 nt dsRNA** | | |
| 24 nt top | 5'- ACAAGCGAAUGAGUCAUUCAUCCU -3' | 2A |
| 25 nt bottom | 5'- AGGAUGAAUGACUCAUUCGCUUGUC -3' | 2A |
| **oligos for testing DCL3 cleavage of blunt-ended 23-24 nt dsRNAs** | | |
| 24 nt top | 5'- AGGAUGAAUGACUCAUUCGCUUGU -3' | 2B |
| 23 nt top | 5'- GGAUGAAUGACUCAUUCGCUUGU -3' | 2B |
| 24 nt bottom | 5'- ACAAGCGAAUGAGUCAUUCAUCCU -3' | 2B |
| **oligos for testing DCL3 cleavage of 23-24 nt dsRNAs with 3' overhangs** | | |
| 24 nt top | 5'- ACAAGCGAAUGAGUCAUUCAUCCU -3' | 2C |
| 24 nt bottom 3' overhangs | 5'- GAUGAAUGACUCAUUCGCUUGUAA -3' | 2C |
| 23 nt bottom 3' overhangs | 5'- GAUGAAUGACUCAUUCGCUUGUA -3' | 2C |
| **oligos for testing DCL3 cleavage of dsRNAs <24 nt** | | |
| 23 nt top | 5'- ACAAGCGAAUGAGUCAUUCAUCC -3' | 2D |
| 23 nt bottom | 5'- GGAUGAAUGACUCAUUCGCUUGU -3' | 2D |
| 22 nt bottom | 5'- GAUGAAUGACUCAUUCGCUUGU -3' | 2D |
| 22 nt top | 5'- ACAAGCGAAUGAGUCAUUCAUC -3' | 2D |
| **oligos for testing RNaseIII domain dicing** | | |
| 26 nt top | 5'- ACAAGCGAAUGAGUCAUUCAUCCUAA -3' | 3B, S2 |
| 27 nt bottom | 5'- UUAGGAUGAAUGACUCAUUCGCUUGUA -3' | 3B |
| 28 nt bottom | 5'- UUAGGAUGAAUGACUCAUUCGCUUGUAG -3' | S2 |
| **oligos for testing DCL3 5' nucleotide preferences** | | |
| 38 nt top 5' A | 5'- ACAAGCGAAUGAGUCAUUCAUCCUAAGUCCAACAUAGU -3' | 4A |

| | | |
|---|---|---|
| 39 nt bottom 5' A | 5'- ACUAUGUUGGACUUAGGAUGAAUGACUCAUUCGCUUGUA -3' | 4A |
| 38 nt top 5' U | 5'- UCAAGCGAAUGAGUCAUUCAUCCUAAGUCCAACAUAGU -3' | 4A |
| 39 nt bottom 5' U | 5’- ACUAUGUUGGACUUAGGAUGAAUGACUCAUUCGCUUGAA -3' | 4A |
| 38 nt top 5' C | 5’- CCAAGCGAAUGAGUCAUUCAUCCUAAGUCCAACAUAGU -3’ | 4A |
| 39 nt bottom 5' C | 5’- ACUAUGUUGGACUUAGGAUGAAUGACUCAUUCGCUUGGA -3’ | 4A |
| 38 nt top 5' G | 5’- GCAAGCGAAUGAGUCAUUCAUCCUAAGUCCAACAUAGU -3’ | 4A |
| 39 nt bottom 5' G | 5’- ACUAUGUUGGACUUAGGAUGAAUGACUCAUUCGCUUGCA -3’ | 4A |
| **oligos for testing DCL3 3' nucleotide preference** | | |
| 38 nt top 5' A | 5'- ACAAGCGAAUGAGUCAUUCAUCCUAAGUCCAACAUAGU -3' | 4B |
| 39 nt bottom 3' A | 5'- ACUAUGUUGGACUUAGGAUGAAUGACUCAUUCGCUUGUA -3' | 4B |
| 39 nt bottom 3' U | 5'- ACUAUGUUGGACUUAGGAUGAAUGACUCAUUCGCUUGUU -3' | 4B |
| 39 nt bottom 3' C | 5'- ACUAUGUUGGACUUAGGAUGAAUGACUCAUUCGCUUGUC -3' | 4B |
| 39 nt bottom 3' G | 5'- ACUAUGUUGGACUUAGGAUGAAUGACUCAUUCGCUUGUG -3' | 4B |
| **oligos for testing DCL3 5' phosphate group preferences** | | |
| 39 nt top | 5'- AACAUGUUGGACUUAGGAUGAAUGACUCAUUCGCUUGUU -3' | 4C |
| 39 nt bottom 5' triphosphate | 5'$^{PPP}$- ACAAGCGAAUGAGUCAUUCAUCCUAAGUCCAACAUGUUU -3' | 4C |
| 39 nt bottom | 5'- ACAAGCGAAUGAGUCAUUCAUCCUAAGUCCAACAUGUUU -3 | 4C |
| **oligos for testing DCL3 ATP-dependency** | | |
| 37 nt top | 5'- ACAAGCGAAUGAGUCAUUCAUCCUAAGUCCAACAUAG -3' | 5 |
| 39 nt bottom | 5'- CUAUGUUGGACUUAGGAUGAAUGACUCAUUCGCUUGUAG -3' | 5 |

**Supplementary Table 2: Oligonucleotides used for Pol IV and RDR2 transcription**

| **Name** | **Sequence** | **Related to Figure(s)** |
|---|---|---|
| **RNA oligo used for Pol IV-RDR2 transcription reaction** | | |
| 16 nt RNA primer | 5'- UGCUUUUUGUCCUGGC -3' | 6B, 6D |
| **DNA oligos used for Pol IV-RDR2 transcription reaction** | | |
| 51 nt T-less template DNA | 5’CAAAAACGAGACAGACAACGAAAGCAGACAGAGAACGCCAG GACCGACACG -3’ | 6B, 6D |
| 28 nt non-template DNA | 5’- GCTGCTTTCGTTGTCTGTCTCGTTTTTG -3’ | 6B, 6D |
| **RNA oligo used for recombinant RDR2 transcription reaction** | | |
| 37 nt template RNA | 5'- UGCGUGCGUCUGCGUCGUUCGUCCUUUGUCCUUCUUC3$'^{\text{dideoxy}}$ | 6C |

**Supplementary Table 3. Reagents and resources used in the study**

| REAGENT or RESOURCE | SOURCE | IDENTIFIER |
|---|---|---|
| Antibodies | | |
| Anti-FLAG® M2 affinity gel | Sigma Aldrich | Cat# A2220 |
| Bacterial and Virus Strains | | |
| Mix & Go Competent Cells, Strain DH5 Alpha | Zymo Research | Cat# T3007 |
| DH10Bac | Invitrogen | Cat# 10361012 |
| DH10EMBacY | Geneva Biotech | N/A |
| Chemicals, Peptides, and Recombinant Proteins | | |
| Plant protease inhibitor cocktail, 100X (in DMSO) | Sigma Aldrich | Cat# P9599 |
| PMSF | Sigma Aldrich | Cat# P7626 |
| GlycoBlue™ | Thermo Fisher | Cat# AM9515 |
| Ribolock RNase Inhibitor | Thermo Fisher | Cat# EO0384 |
| RNase Inhibitor Murine | NEB | Cat# M0314 |
| RNase H | NEB | Cat# M0297S |
| RNase I | Promega | Cat# 4261 |
| S1 nuclease | Promega | Cat# M5761 |
| S1 nuclease | Thermo Fisher | Cat# EN0321 |
| Vaccinia capping system | NEB | Cat# M2080S |
| Proteinase K, RNA grade | Invitrogen | Cat# 25530049 |
| T4 Polynucleotide Kinase | NEB | Cat# M0201S |
| Finale herbicide | Bayer | CAS# 77182822 |
| Adenosine 5'-triphosphate magnesium salt | Sigma Aldrich | Cat# A9187 |
| Adenosine 5′-[γ-thio]triphosphate tetralithium salt | Sigma Aldrich | Cat# A1388 |
| Set of rATP, rUTP, rCTP, rGTP | Promega | Cat# E6000 |
| Grace's insect cell media, supplemented | Thermo Fisher | Cat# 11605-102 |
| Fetal Bovine Serum, certified, United States | Thermo Fisher | Cat# 16000069 |
| Express Five™ SFM | Thermo Fisher | Cat# 10486025 |
| RNA Loading Dye (2x) | NEB | Cat# B0363S |
| 2x TBE-Urea Sample Buffer | Invitrogen | Cat# LC6876 |

| | | |
|---|---|---|
| IPEGAL® CA-630 | Sigma Aldrich | Cat# I8896 |
| SYBR® Gold Nucleic Acid Gel Stain (10,000x) | Invitrogen | Cat# S11494 |
| 3x (DYKDDDDK) peptide | APExBio | Cat# A6001 |
| Critical Commercial Assays | | |
| High Pure Viral RNA kit | Roche | Cat# 11858882001 |
| Raw and analyzed data | This paper | |
| Experimental Models: Cell Lines | | |
| Sf9 in Grace's Medium | Thermo Fisher | Cat# B82501 |
| High Five™ Cells in Express Five Medium | Thermo Fisher | Cat# B85502 |
| Experimental Models: Organisms/Strains | | |
| *Arabidopsis thaliana* strain Col-0 | N/A | N/A |
| NRPD1-FLAG in Col-0 | Haag et al., 2012 | N/A |
| Oligonucleotides | | |
| For DCL3-only dicing assays (Figures 1-5) please see Table S1 | | |
| For Pol IV RDR2 transcription reactions (Figure 6) please see Table S2 | | |
| Recombinant DNA | | |
| pUC57-DCL3 (Codon optimized for insect cells) | Singh et al., 2019 | (Genscript) N/A |
| pFastBac™ HT B-DCL3 | Singh et al., 2019 | N/A |
| pFastBac™ HT B-DCL3_E1146Q_E1329Q | This paper | N/A |
| pFastBac™ HT B-DCL3_E1146Q | This paper | N/A |
| pFastBac™ HT B-DCL3_E1329Q | This paper | N/A |
| Software and Algorithms | | |
| Image Lab v6.0.1 | Bio-Rad | Cat # 12012931 |

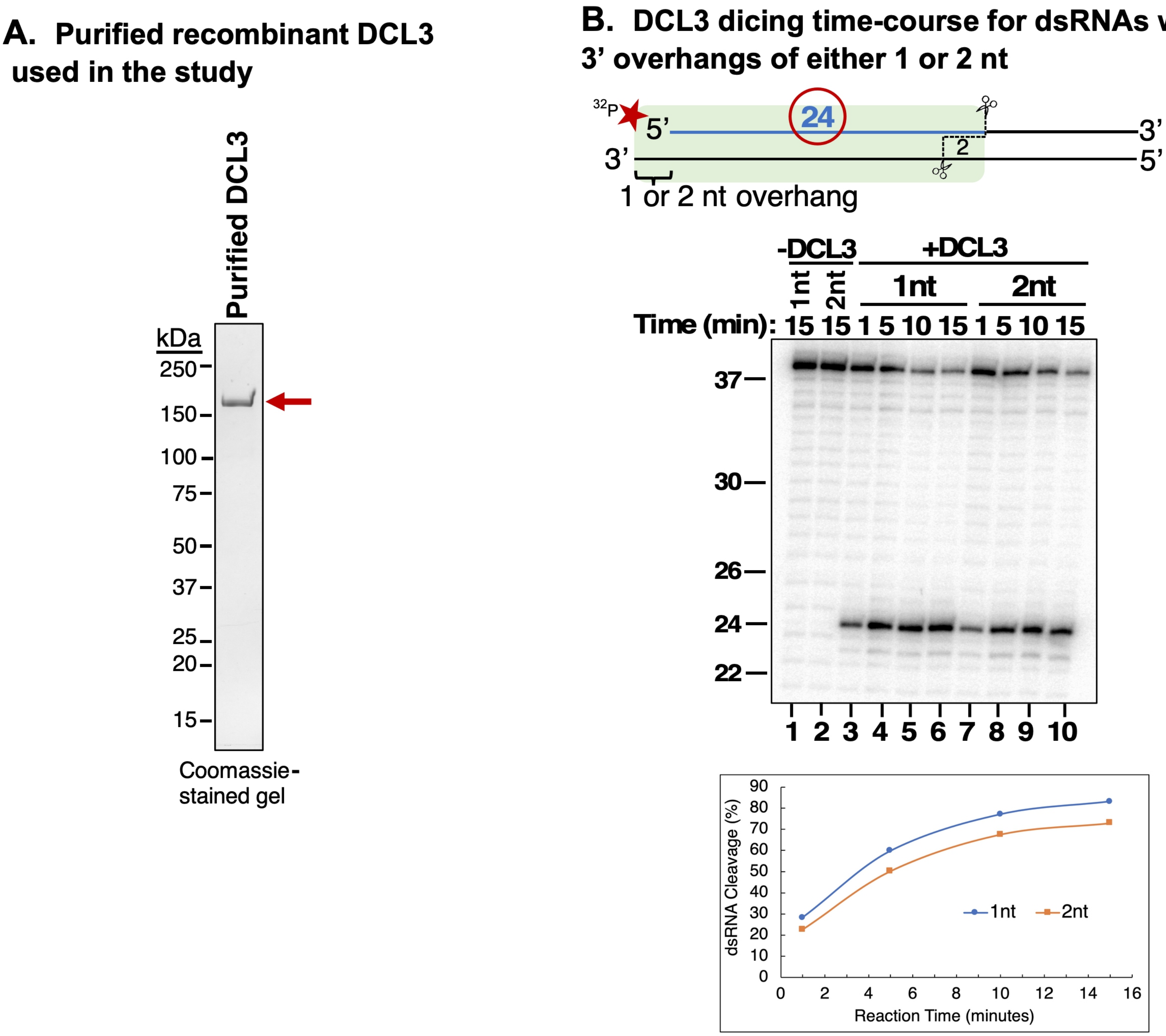

**Figure 1_Figure supplement 1.**
Affinity Purification of recombinant DCL3.
A. Recombinant wild-type (WT) DCL3 bearing a C-terminal FLAG tag was produced in insect cells and affinity purified using anti-FLAG resin. The purified protein was analyzed by SDS-PAGE and Coomassie Blue staining.
B. Time course comparison of DCL3 dicing of substrates with either 1 nt or 2 nt 3' overhangs. An end-labeled 37 nt top strand was annealed to 38 or 39 nt bottom strands to form dsRNA substrates with either 1 or 2 nt 3' overhangs. Resulting dsRNAs (50 nM) were then incubated with 25 nM of affinity purified recombinant DCL3 for 1, 5, 10 or 15 minutes. RNAs were then resolved by denaturing polyacrylamide gel electrophoresis (PAGE) and visualized using SYBR Gold staining. Lanes 1 and 2 are controls that contain the different dsRNAs but no protein.

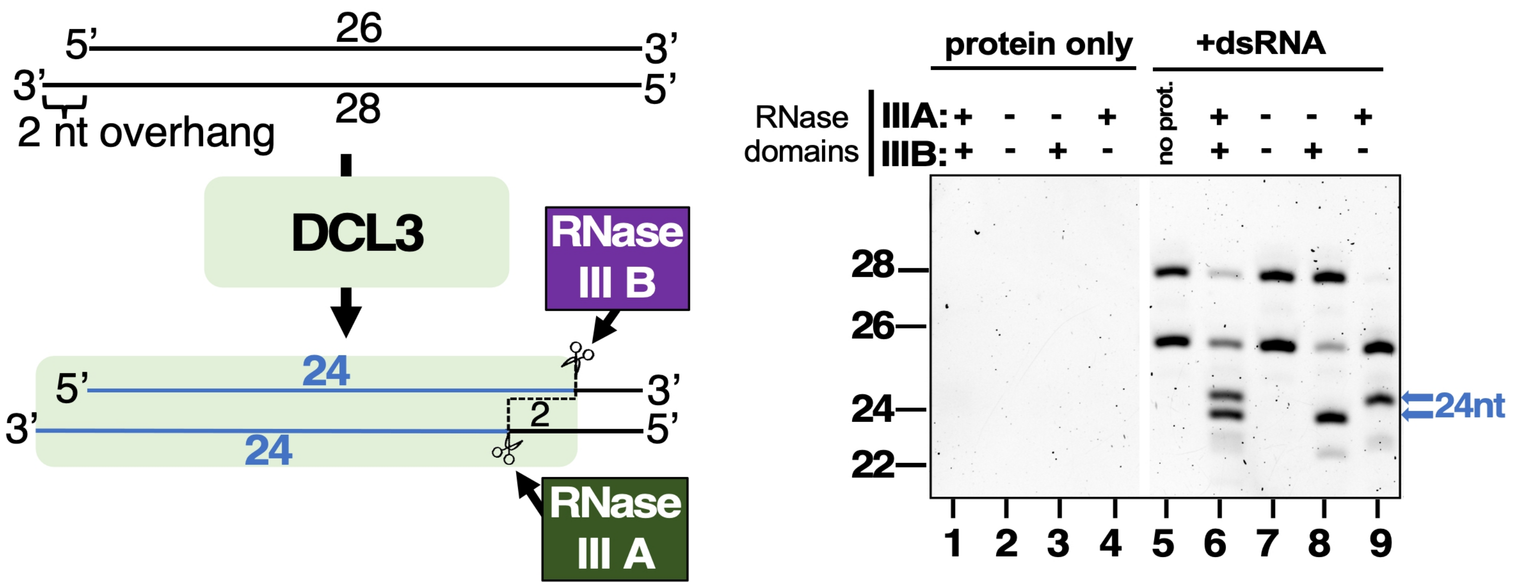

**Figure 3_Figure supplement 1**

Strand cutting specificities of DCL3's two RNase III domains tested using a dsRNA substrate with a 2 nt 3' overhang.

This experiment is similar to that of Figure 3 except that the dsRNA substrate has a 2 nt overhang rather than a 1 nt overhang. In this case a dsRNA formed by annealing 26 and 28 nt RNAs was subjected to dicing using wild-type DCL3 or the E1146Q and/or E1329Q mutant versions of DCL3. Lanes 1-4 are DCL3-only controls and lane 5 is a RNA-only control. DCL3 with wild-type RNAse III domains A and B (denoted as +, +) was tested in lanes 1 and 6. DCL3 mutants with both RNase III domains mutated (denoted as -, -) was tested in lanes 2 and 7. Mutants with only wild-type RNase III domain were tested in lanes 3,4,8 and 9).
